# The menstrual cycle is a primary contributor to cyclic variation in women’s mood, behavior, and vital signs

**DOI:** 10.1101/583153

**Authors:** Emma Pierson, Tim Althoff, Daniel Thomas, Paula Hillard, Jure Leskovec

## Abstract

Female mood, behavior, and vital signs exhibit cycles which fundamentally affect health and happiness. However, it is unclear which dimensions of mood, behavior, and vital signs vary cyclically, how cycles at different timescales compare in magnitude, and how cycles vary across countries. Here we separate female mood, behavior, and vital signs into four simultaneous cycles – daily, weekly, seasonal, and menstrual. We analyze nine mood dimensions, three behavior dimensions, and three vital signs using a dataset of 241 million observations from 3.3 million women in 109 countries. We find that the menstrual cycle is a primary contributor to cyclic variation: it is the cycle with the largest amplitude for all three vital signs, sexual activity, and 7 out of 9 dimensions of mood. The amplitude of the menstrual happiness cycle is 1.4x the amplitude of the daily cycle, 3.3x the amplitude of the weekly cycle, 2.3x the amplitude of the seasonal cycle, and 1.7x the Christmas increase in happiness. Menstrual cycle effects are directionally consistent across countries, demonstrating that they occur across cultures. Overall, our results demonstrate the primacy of the menstrual cycle and necessitate better accounting for it in clinical data and practice.

---

Female mood, behavior, and vital signs exhibit cycles over multiple timescales – for example, the day, the week, and the menstrual cycle – and these cycles have important health consequences. Seasonal cycles are implicated in mood disorders (1), circadian cycles in sleep (2) and obesity (3), and the menstrual cycle in fertility (4), schizophrenia (5), and cancer (6). Previous work has uncovered evidence of cycles on four timescales: daily, weekly, seasonal, and menstrual (1–11).

However, three fundamental questions about these cycles remain unresolved. First, it is not clear what dimensions actually cycle because previous studies of daily (7), seasonal (10), and menstrual cycles (9, 11) have yielded conflicting results. For example, previous studies of peaks in negative mood have disagreed about what time of day they occur (7), what time of year they occur (10), where in the menstrual cycle they occur (9), and even whether they occur at all. Second, while there is evidence of daily, weekly, seasonal, and menstrual cycles, no study has compared these four cycles simultaneously, and thus their relative magnitudes are unknown. Third, because many previous datasets have been small-scale and country-specific, they have been unable to study how factors like cultural background and age affect cycle dynamics, and it is unclear to what extent their conclusions generalize (7). Uncertainty over how cyclic patterns generalize across cultures has clinical implications: for example, it was a primary argument against including premenstrual dysphoric disorder in the Diagnostic and Statistical Manual of Mental Disorders (DSM) (12). Recently, analyses of large-scale datasets — for example, sentiment analysis of Twitter (7) — have studied cycles across broader populations. However, these datasets have lacked information on menstrual cycles, so while they have examined daily, weekly, and seasonal cycles, they have been unable to simultaneously study the menstrual cycle.

We provide the first decomposition of women’s mood, behavior, and vital signs into simultaneous daily, weekly, seasonal, and menstrual cycles. We use an international dataset of 241 million observations from 3.3 million reproductive age women who use the women’s health mobile-tracking application Clue by BioWink GmbH, which allows women across more than 100 countries to prospectively track more than 100 features (Figure 1, Tables S1 and S2). Such data from women’s health tracking apps has only recently become available on a large scale as the apps have grown in popularity; a critical advantage these datasets offer is information on menstrual cycle starts, enabling comparison of the menstrual cycle to daily, weekly, and seasonal cycles for the first time. The Clue app has been rated the most accurate menstrual cycle tracking app by gynecologists (13) and previously used in studies which show that it replicates known biological findings (14–17). Using the Clue dataset, we analyze menstrual, daily, weekly, and seasonal cycles in nine dimensions of mood, three vital signs — resting heart rate (RHR), basal body temperature (BBT), and weight — and three dimensions of behavior — sleep, exercise, and sexual activity (Tables S2 and S3). We choose these dimensions because they are fundamental to human health and well-being, previous research has suggested they may exhibit cycles across multiple timescales, and they are logged by large numbers of people in our dataset.

**Figure 1:**
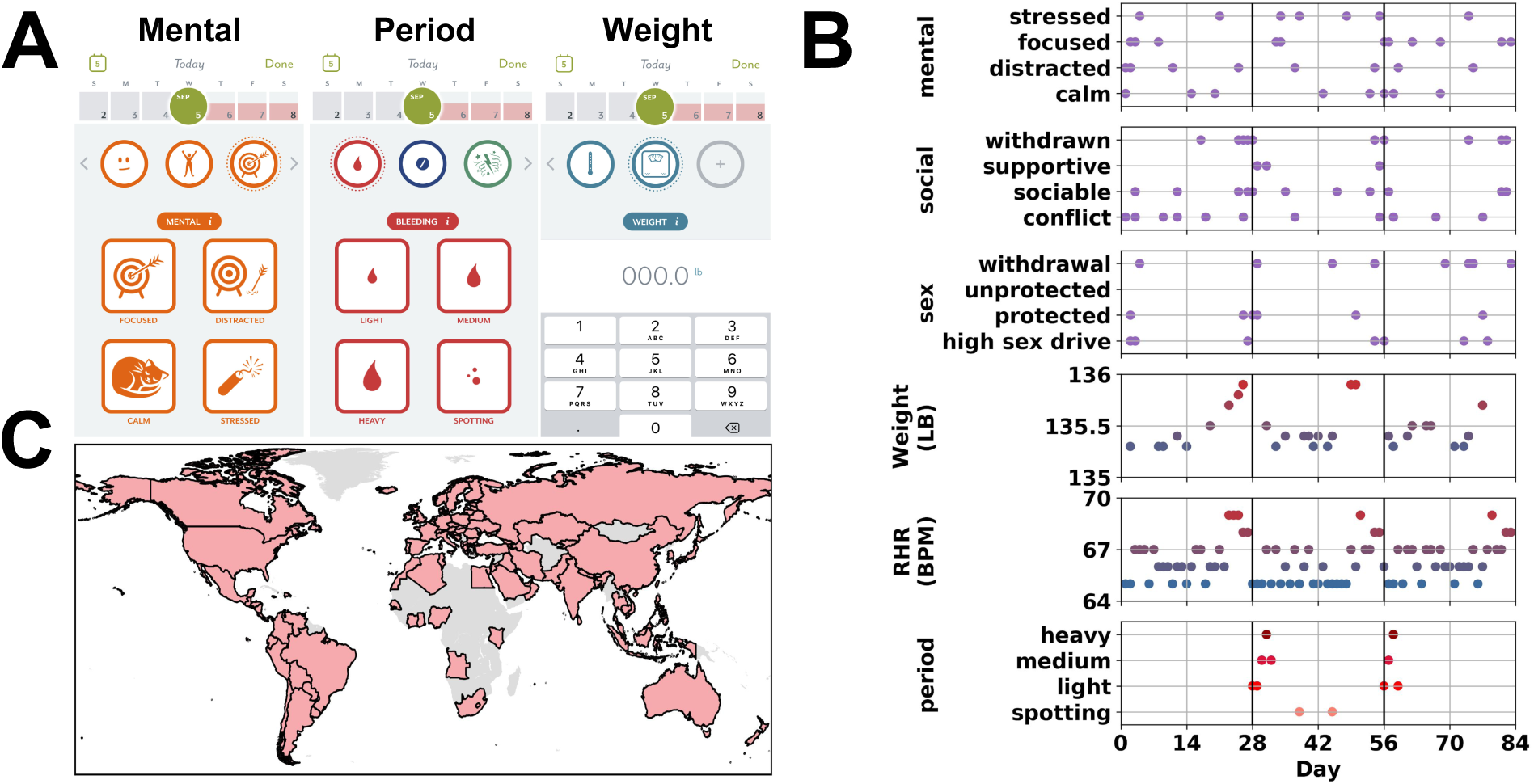
Overview of the dataset. **A**: Three screenshots from the women’s health mobile app which was used to collect data, illustrating how women can enter logs about their mental state, period starts, and weight. Mental and period logs are binary, and can be entered by tapping a square on the screen; weight is continuous and entered as a numerical value. **B**: illustrative simulated data for a single person; the horizontal axis is time, and the vertical black lines denote inferred menstrual cycle starts (Methods). Each dot represents the person logging one feature on one day; only a subset of the features that can be logged are shown. The top three subplots illustrate the logging of binary features; the next two subplots illustrate weight and resting heart rate (RHR), which are continuous; period logging data is shown in the bottom subplot. **C**: The 109 countries with at least 100 people and 1,000 observations in the dataset.

We first illustrate that failing to account for the menstrual cycle substantially understates individual cyclic variation (Figure 2). Many previous studies of cyclic variation analyze a daily signal averaged across the whole population (7, 8) and we thus begin with this analysis for the happy-sad dimension of mood (Figure 2a)^1^. This reveals some cycles, like weekly cycles, as well as outliers like Christmas (when happiness increases) and the day after the 2016 US Election (when happiness decreases). But averaging across the whole population conceals the menstrual cycle because menstrual cycles do not begin on the same day for every woman, and so menstrual variation is averaged out when all women are combined. To reveal this variation, we leverage the fact that our dataset contains the dates when menstrual cycles begin (*i.e*., the dates when the period begins; Methods) for each person. This allows us to break happy/sad mood into menstrual, daily, weekly, and seasonal cycles by running a linear regression of happy/sad mood on categorical variables for day relative to period start, hour of day, day of week, and month of year (Methods, Figure 2b). We find that the menstrual cycle is larger in amplitude (as measured by the cycle maximum minus its minimum) than the other three cycles for 7 out of 9 dimensions of mood, sexual behavior, and all three vital signs, making it a primary contributor to cyclic variation (Figure 2c). For happy-sad mood, the amplitude of the menstrual cycle (5.5%) is 1.4x the amplitude of the daily cycle, 3.3x the amplitude of the weekly cycle, and 2.3x the amplitude of the seasonal cycle. Importantly, the amplitude of the menstrual cycle is also significant relative to the overall mean: on average, 25% of happy/sad logs are sad, so the 5.5% menstrual cycle amplitude is a 22% relative change in the probability of logging sadness. The amplitude of the menstrual cycle is also significant relative to the effects of outlier events: it is about 1.7x the Christmas effect and 0.6x the 2016 US Election effect. Consequently, if the menstrual variation could be observed in the population signal (Figure 2a, red line) it would substantially increase the cyclic variability. Studies of cyclic variation in mood, behavior, and vital signs will be more accurate if menstrual cycles are accounted for.

**Figure 2:**
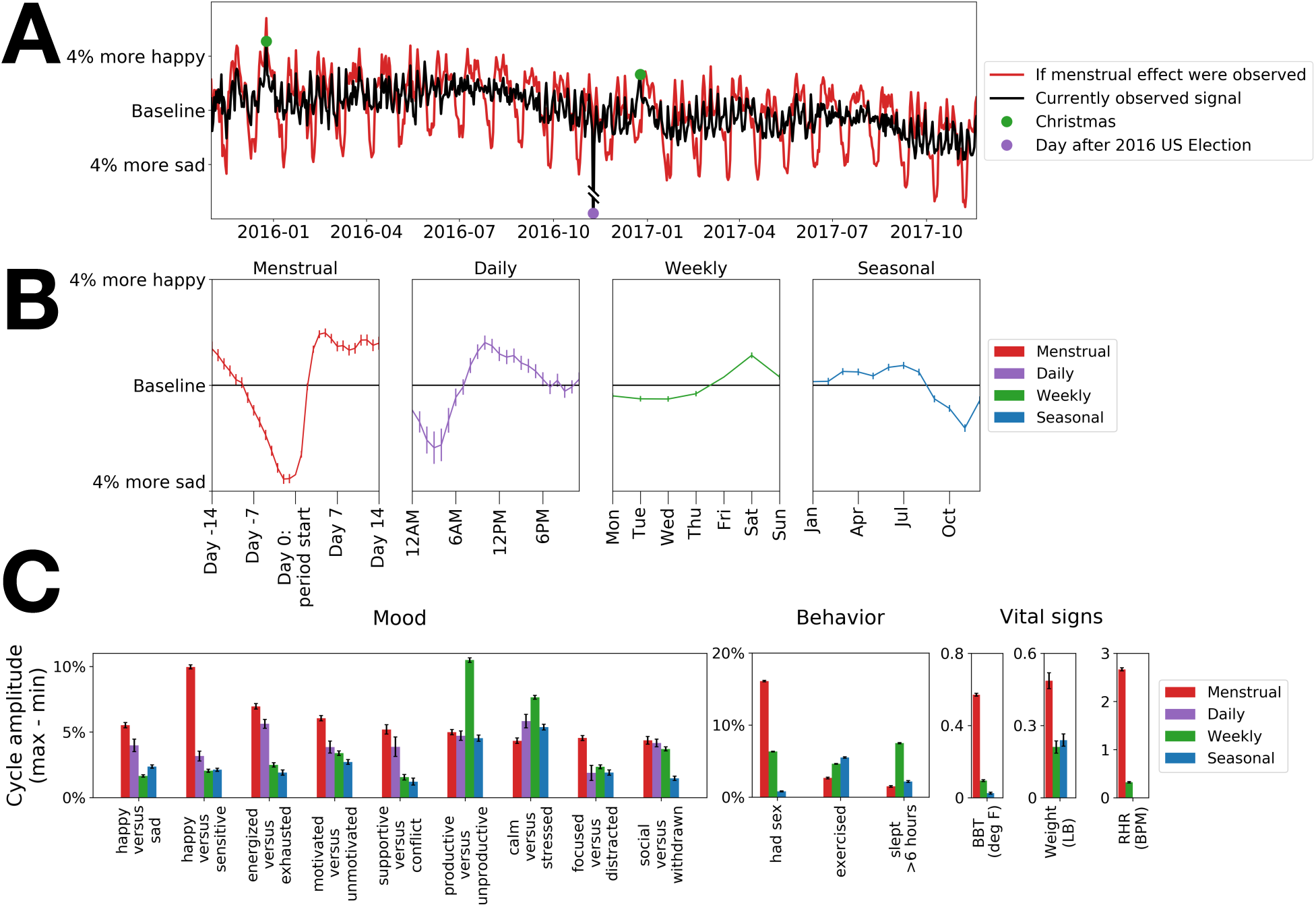
Decomposition of female mood, behavior, and vital signs into four simultaneous cycles reveals that the menstrual cycle is a primary contributor to cyclic variation. **A**: Averaging out the menstrual cycle conceals variation in happy-sad mood. The black line plots the fraction of happy-sad logs which are happy by day, after subtracting the baseline (individual mean) for each person to reveal within-individual variation. Weekly cycles are apparent, as well as outlier events like Christmas and the day after the 2016 US Election, but this view is also limited: menstrual cycles cannot be seen because cycles do not begin on the same day for each woman, and so are averaged out. The red line plots what the black line would look like if the menstrual cycle could be observed (Methods), providing a more accurate picture of individual variability. The amplitude of the menstrual cycle is 1.7x the Christmas effect and 0.6x the 2016 US Election effect. (To focus on the predominant range of variation, the vertical axis truncates the US Election outlier, which actually appears 10.3% below baseline.) **B**: Happy-sad mood can be broken down into menstrual, daily, weekly, and seasonal cycles (Methods). **C**: amplitude of cycles for all mood, behavior, and vital sign dimensions. Menstrual cycles (red bars) have the largest amplitude for 7 out of 9 dimensions of mood, sexual behavior, and all three vital signs: basal body temperature (BBT), weight, and resting heart rate (RHR). The weekly cycle (green) is the largest cycle for productivity, stress, and sleep, the daily cycle (purple) is prominent for most mood dimensions, and the seasonal cycle (blue) is relatively small in magnitude. For behavior and vital signs, no hourly information is available, so the daily cycle is not shown; for RHR, less than a year of data is available, so the seasonal cycle is not shown.

We emphasize that we analyze variation *within* people, as opposed to *between* people (7, 8, 11). For example, if a given person weighs twenty pounds less than the population mean, most of that variation is likely due not to cyclic variation but to between-person variation, since the menstrual weight cycle amplitude is roughly half a pound (Figure S3). Our analysis implies that the menstrual cycle is a primary contributor to within-individual cyclic variation.

We next analyze the four-cycle decompositions of all 15 dimensions in our analysis, revealing intriguing variation. While the menstrual cycle is greater in amplitude than the other three cycles for 7 out of 9 dimensions of mood (happy/sad, happy/sensitive, energized/exhausted, focused/distracted, motivated/unmotivated, sociable/withdrawn, supportive social/conflict social), the daily cycle is also prominent, with mood becoming more negative between midnight and 6 AM (Figure S1). While seasonal and weekly cycles in mood are generally smaller than the daily and menstrual cycles, they are large for the calm/stressed and productive/unproductive dimensions: calm peaks during the summer and end-of-year (likely due to school holidays, since a large fraction of the population in the dataset is school-age) and on weekends, and productivity declines. These results speak to the importance of modeling multiple dimensions of mood, which may show different cyclic patterns. For behavior dimensions, the menstrual cycle is the largest cycle for sexual activity, with a large decrease in logged sexual activity immediately after the period begins; sexual activity also increases on the weekends (Figure S2). For sleep, the weekly cycle is most prominent, with an increase in sleep on the weekends; for exercise, both weekly and seasonal cycles are prominent, with decreases in exercise on the weekends and during the winter (Figure S2). The menstrual cycle is the largest cycle for all three vital signs — basal body temperature (BBT), resting heart rate (RHR), and weight (Figure S3).

We confirm that the amplitudes and cyclic patterns we observe remain stable under a number of robustness checks (Methods). First, we repeat our analysis under alternate statistical models, including two other cycle decomposition methods; alternate parameterizations of the seasonal cycle; and alternate parameterizations of the menstrual cycle. As a second robustness check, we repeat our main analyses across subsets of the dataset broken down by app usage variables and demographics. While, as expected, we observe some variation across subsets of the population, we find that the ordering of cycle amplitudes remains generally stable (Figure S5) and, importantly, the prominence of the menstrual cycle is not driven by a particular subgroup. This shows that while the population in our dataset is likely non-representative, the prominence of the menstrual cycle persists across subpopulations, and our conclusions are more likely to generalize.

Having established the importance of the menstrual cycle, we next examine how it varies across countries. Because most previous studies of the menstrual cycle have been small-scale and country-specific, it has been unclear to what extent menstrual effects persist across countries. This uncertainty has clinical implications: for example, whether premenstrual mood disorder should be included in the DSM-V has been disputed on the grounds that it might be culturally specific (12). To examine how menstrual effects vary across countries, we define the *near-period effect* for a dimension as the difference in the mean value of the dimension when an individual is near their period start and when they are not (Methods). Contrary to previous concerns that near-period effects are culturally specific, we find they are directionally consistent across countries. For example, the near-period decrease in happiness occurs across all 87 countries we examine (Figure 3). The other large near-period effects in mood, sexual behavior, and vital signs also remain directionally consistent across countries (Figure S6). We confirmed that our estimates of country-specific period effects remained consistent (Methods, Figure S7) when we controlled for demographic covariates, behavior covariates, and app usage covariates.

**Figure 3:**
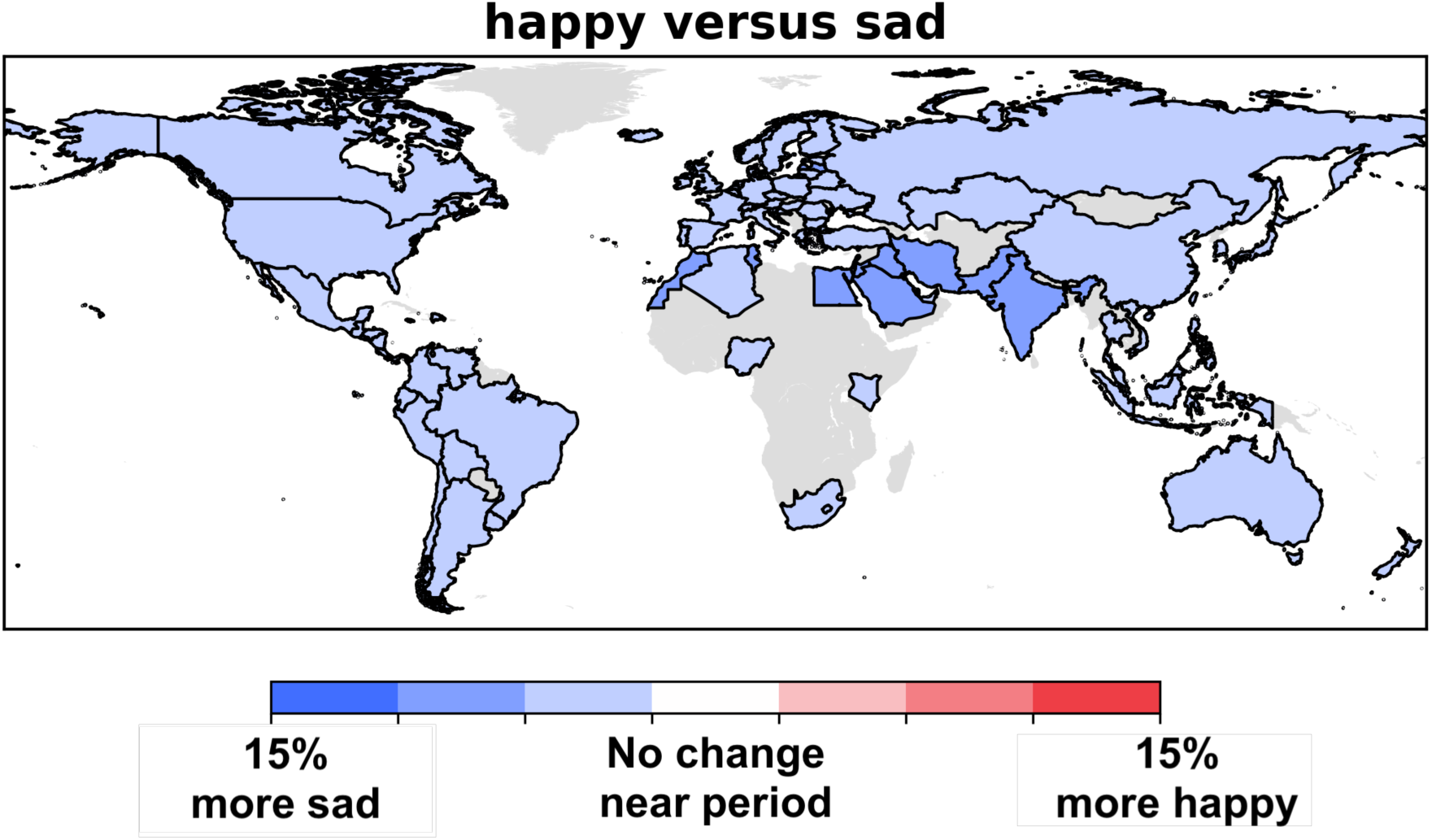
Near-period mood effects are directionally consistent across countries, demonstrating that they occur across cultures. The *near-period effect* for a dimension is the difference between the mean value of the dimension when an individual is near their period start and when they are not. All countries show an increase in negative mood near the start of the period. Color bins are equally sized increments: blues indicate a negative change in mood near the period, the central white bin is centered around no change near the period, and reds indicate a positive change in mood near the period. Countries with fewer than 1,000 observations and 100 individuals in the dataset are shown in grey. Near-period effects in the other dimensions are directionally consistent as well (Figure S6).

We next examine how near-period effects vary by age (Figure 4). Menstrual cycle dynamics are known to change with age (18–22) and understanding normal aging-related changes is important for characterizing healthy menstrual patterns (20); however, it has not been possible to study aging trends in all the dimensions we consider on the scale of our dataset. The near-period negative mood effect increases with age (from 3.6% in 15-20 year olds to 5.4% in 30-35 year olds, a relative increase of 51%), consistent with prior reports that premenstrual dysphoria can increase during the late reproductive years (20). We also observe age trends in all three vital signs: with increasing age, the near-period effects for resting heart rate and weight decrease in magnitude, while the near-period effect for BBT increases. The increase in the near-period BBT effect with age is consistent with the fact that the fraction of cycles in which ovulation occurs increases with age (21), and BBT rises at ovulation (23). While all these trends are robust to the inclusion of controls for demographics, behavior, and app usage (Figure S8), it is possible that unobserved heterogeneity also contributes to the age trends: for example, the age trend in BBT may also be driven in part by the fact that BBT is difficult to measure properly, and young women may be less skilled at it^2^. Because our analysis is cross-sectional (since our median followup time, of only 1.47 years for each individual, is too short to allow longitudinal analysis), longer followup on large longitudinal datasets should further investigate the trends we observe.

**Figure 4:**
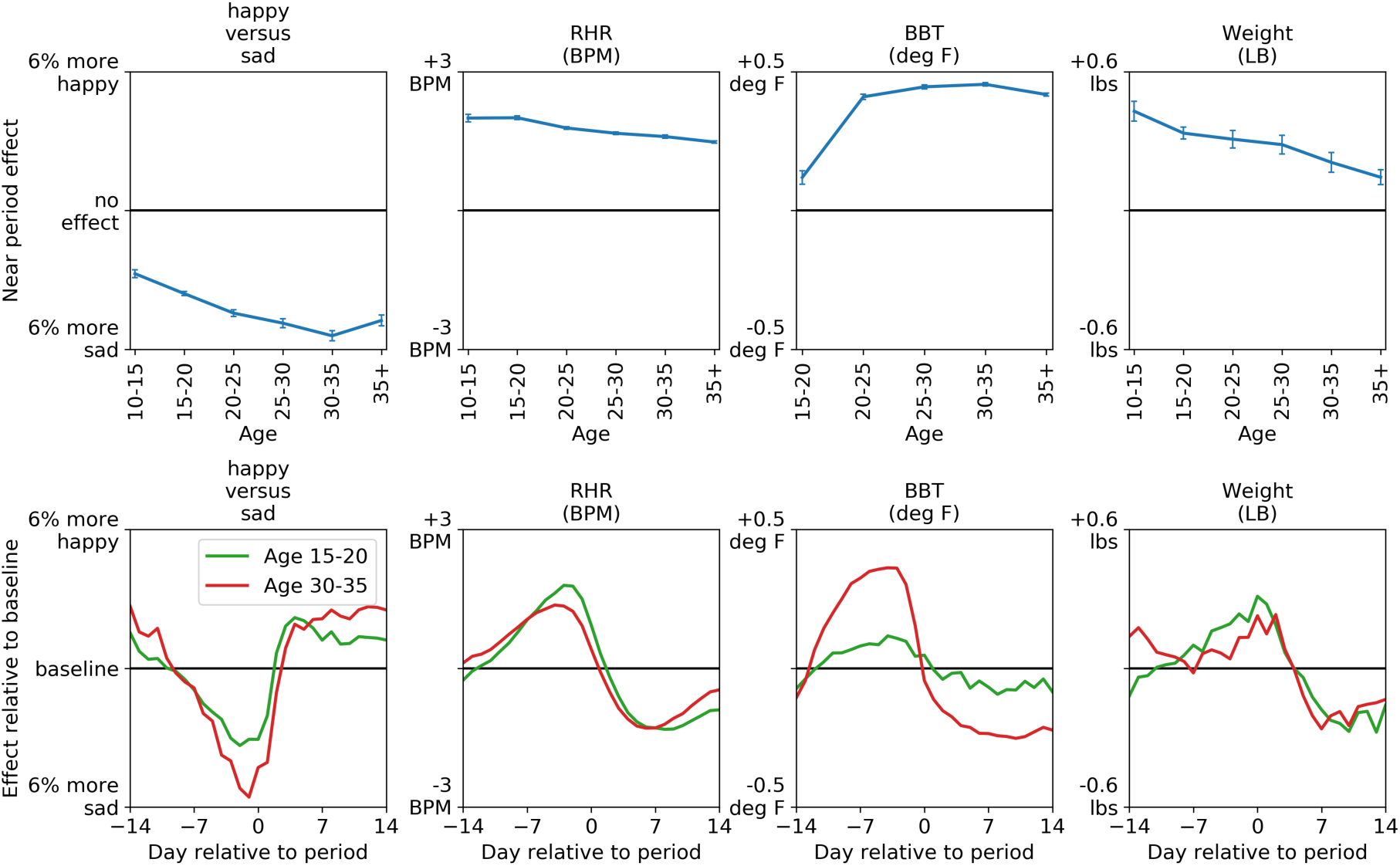
Near-period effects change with age for happy-sad mood, resting heart rate (RHR), basal body temperature (BBT), and weight. The top row of plots show near-period effects (vertical axis) as a function of age (horizontal axis). The near-period effect for happy-sad mood grows larger with age (from 3.6% in 15-20 year olds to 5.4% in 30-35 year olds, a relative increase of 51%). The near-period change in RHR grows smaller with age (from 2.0 BPM in 15-20 year olds to 1.6 BPM in 30-35 year olds, a relative decrease of 20%). The near-period effect for BBT increases with age (from 0.12 deg F in 15-20 year olds to 0.45 deg F in 30-35 year olds, a relative increase of 281%). The near-period effect for weight decreases with age (from 0.33 lbs in 15-20 year olds to 0.21 lbs in 30-35 year olds, a relative decrease of 38%). All 4 plots show statistically significant changes with age, with errorbars denoting 95% confidence intervals (Methods); estimates of age effects are robust to inclusion of controls for demographics, behavior, and app usage (Figure S8). The bottom row of plots show the change in each dimension over the course of the menstrual cycle for 15-20 year olds (green line) and 30-35 year olds (red line). (All age groups have at least 1,000 observations and 100 individuals in the dataset; for BBT, the age group 10-15 is not shown because women in this age group log BBT too infrequently.)

Our dataset has limitations. First, the population in our dataset — smartphone users using a women’s health app — is not representative of the global population: it is potentially biased towards individuals of higher socioeconomic status or who are particularly interested in women’s health. This raises the question of whether our findings generalize. As we discuss in more detail in the Methods section, several lines of evidence mitigate this concern: a) menstrual cycle apps are increasingly widely used (24), offering a data source which is arguably more representative than previous studies using small and non-representative populations (25); b) previous analyses of this dataset (14–17) have reproduced numerous results consistent with previous findings, and we further confirm a number of previous results in our present analysis (Methods); and c) the menstrual cycle remains prominent across subsets of the dataset, suggesting that this finding is not driven by a single non-representative population. A second limitation of our dataset is that it relies on self-reported data, which may not always be reliable, particularly for dimensions like BBT which require skill to measure accurately. Again, multiple lines of evidence mitigate this concern: first, a review of more than a hundred menstrual cycle tracking apps rated the Clue app as the best on the basis of accuracy, comprehensiveness, and functionality (13); second, we apply numerous quality control filters to increase the accuracy of logged data. The medical community is already making use of menstrual app tracking data (14–17, 25), suggesting that neither of these limitations precludes its usefulness.

Our finding that the menstrual cycle is a primary contributor to cyclic variation has implications in three areas: medical data collection, clinical practice, and the broader cultural framing of the menstrual cycle. First, in terms of data collection, menstrual cycle information is often lacking from medical records (25) and global health data (26); health-tracking apps have similarly been slow to incorporate menstrual tracking (27). Without collecting menstrual cycle data, it is impossible to account for and study this fundamental source of variation. Second, in terms of clinical practice, clinicians do not always consider menstrual health (28), in spite of the fact that the menstrual cycle is considered a vital sign which can be used to monitor numerous health conditions (22, 29). Our findings emphasize the role of the menstrual cycle not just in disease, but in many dimensions of variation relevant to patients’ well-being. Third, the menstrual cycle has been poorly understood and even stigmatized in popular culture, undermining women’s health (25, 29). Our analysis shows that the menstrual cycle, as a primary contributor to cyclic variation in women’s mood, behavior, and vital signs, must be normalized and understood just as daily, weekly, and seasonal cycles are.

## Acknowledgments

The authors thank Pang Wei Koh, Shengwu Li, Adam Mastrioanni, Maria Mateen, Leah Pierson, Rachel Pierson, Nat Roth, Camilo Ruiz, Rok Sosic, Christopher Yau, and Marinka Zitnik for helpful comments, and Hertz and NDSEG Fellowships, the Stanford Data Science Initiative, the Chan Zuckerberg Biohub, and DARPA SIFT for financial support.

## Footnotes

1 To avoid falsely attenuating seasonal cycles, we filter for individuals in the Northern hemisphere who log in at least twelve unique months (Methods).

2 We mitigate this by filtering for regular BBT loggers (Methods).

