## Supplementary material for "The menstrual cycle is a primary contributor to cyclic variation in women’s mood, behavior, and vital signs": Materials and Methods, Supplementary Figures and Tables

### Supplementary Information

1.8 Reliability of self-reported data . . . . .

**2 Supplementary Tables and Figures**

### 1 Materials and Methods

#### 1.1 Clue app interface

In Figure 1 we provide screenshots from the Clue app interface to illustrate how people using the app log symptoms. Related categories of logs appear on the same screen: for example, all period log types (spotting, light, medium, and heavy bleeding) appear on the same screen. All log types except for weight, basal body temperature, and resting heart rate are binary, and can be logged simply by tapping the screen. Each binary log can be entered only once per day. People can backfill information (*i.e.*, enter logs for previous days) by tapping the date at the top of the screen.

#### 1.2 Data processing

##### 1.2.1 Definition of menstrual cycle starts and date relative to period

Consistent with the Clue app, we define a person to be on their period when they log light, medium, or heavy bleeding. We define a *period start* as a start of bleeding when the person has recorded no bleeding for at least 7 days: so if the person records no bleeding in April, and bleeding on May 1, 2, 3, and 29, the period starts would be May 1 and May 29. We remove the first logged period start for every person because some people "backfill" their first period start, potentially creating unreliable dates. We confirm that the distribution of period lengths is consistent with the previous literature (Figure S9). Given the period starts for each person, we define the person's *day relative to period start* on each day $d$ as $d - p$ (where $p$ is the nearest period start as measured by absolute difference in days). So if the closest period start occurred on May 1, the day relative to period on April 30 would be -1, on May 1 would be 0, and on May 2 would be 1. As discussed below in Section 1.5, we confirm that we observe similar cycle trajectories when we compute day relative to period start using only previous period starts or only subsequent period starts (so day relative to period start is always non-negative or always non-positive, respectively).

###### 1.2.2 Location information and privacy

Our dataset contains individuals from the 25,000 cities with the most Clue users, covering 97% of the Clue population. Each city was mapped to its latitude (rounded to the nearest 5 degrees), country, and timezone; Clue removed latitude-timezone pairs with fewer than 1,000 people to protect user privacy, and provided us with the rounded latitude, country, and timezone for each person. We use each person's timezone to compute the local time for each log, and use the person's local time in all analyses. All data is fully de-identified, all Clue users consent to use of data for research purposes, and analysis was determined to be exempt from review by the Stanford Institutional Review Board.

###### 1.2.3 Data filtering

For all dimensions, we filter for logs on dates between November 1, 2015 (since on earlier dates, not all features were available to be logged on the Clue app (*30*), rendering the data incomparable), and November 20, 2017 (when our dataset ends). We also filter out logs that are more than 20 days from a period start, since we will be unable to match these reliably to a period. In our primary analysis, we analyze people who log in at least twelve unique months, whom we term *long-term loggers*. We do this to ensure that we have long enough timespans to estimate seasonal cycles and that people log reliably, mitigating missing data concerns. We verify that our estimates of cycle amplitudes do not change substantially when we examine the remainder of the population (Figure S5).

Because seasonal cycles could potentially be reversed or otherwise altered in the Southern and Northern hemispheres, in our analyses which compare to seasonal cycles (*e.g.*, Figure 2) we analyze only individuals in the Northern hemisphere to avoid falsely attenuating the seasonal cycle. As described in Section 1.5, we verify that cycle amplitudes remain stable when we substratify by latitude subgroups (*e.g.*, individuals between 25° S and the equator). In all analyses which focus on the menstrual cycle and do not consider seasonal cycles (*e.g.*, Figures 3 and 4), we analyze all individuals and do not filter for the Northern hemisphere.

While some previous analyses of the menstrual cycle (*18*) (*14, 32*) have studied only women who are not taking hormonal birth control (and are thus experiencing their natural hormone cycle), we do not apply this filter for two reasons. First, women in our dataset do not reliably log hormonal birth control information, so a hormonal birth control filter would be highly imperfect. Second, the goal of our analysis is to describe cycles in the general population of women, not to describe the effect of the natural hormone cycle specifically. Because hormonal birth control can reduce premenstrual effects (*33*), the menstrual cycle amplitudes we observe might be larger in women who were not taking birth control than in the general population we study.

###### 1.2.4 Binary dimensions: Mood and behavior

In order to ensure we have hourly information for logs and that logs are *prospective* (as opposed to logged weeks after the fact, potentially rendering memory unreliable), we filter for logs for which we have time information on when the logging session started and ended. We remove a very small fraction ($< 0.1\%$) of logs where the session lasts more than an hour, since this renders time information unreliable. To ensure logs are prospective, we remove logs where the person enters a date for the log which is not the same day as the session start.

For each dimension of mood and behavior, we must define a way of converting the categorical data that users log (*e.g.*, tapping “calm” or “happy” on the app screen) to a numerical value on which we can perform quantitative analysis. To define this value for each mood dimension, we pair each mood with an opposing mood: for example, happy with sad, or calm with stressed. The value for a given log is 1 if the person logs the first mood and 0 if they log the opposite mood: for example, 1 if the person logs happy, and 0 if they log sad. (We examine all mood dimensions except for PMS, because it is obviously menstrual-cycle-specific, and high/low energy, because it is redundant with the energized/exhausted dimension we analyze). Thus, the average value of the happy/sad mood dimension can be interpreted as the "fraction of happy/sad logs which are happy". We define mood dimensions in this way to control for the fact that a person's probability of logging fluctuates considerably over time of day and point in the menstrual cycle. For example, simply examining whether a person logs a mood, without normalizing in some way, would show large spikes in all emotions near the menstrual cycle start, due simply to the fact that many women use the app primarily to track menstrual cycle starts and thus are more likely to log other features then as well[3].

We now describe how we map from categorical log data to numerical values for each of the three behavior dimensions: sleep, sex, and exercise. For sleep, the app offers users four logging categories: 0-3 hours, 3-6 hours, 6-9 hours, and more than 9 hours. We define the value of the sleep dimension as 1 if the person logged sleeping 6-9 hours or more than 9 hours, and 0 otherwise. For the sex and exercise behavior dimensions, we use logs of any type as a normalizer: that is, the sex and exercise dimensions are 1 on a day if a person logged that behavior on that day, and 0 if they logged only something other than that behavior. (For the sex dimension, we include only protected, unprotected, and withdrawal sex, and do not include the "high sex drive" feature which the app also allows women to log.) To ensure that people are

[3]Pairing emotions with their opposites is consistent with previous conceptualizations of emotion (*34, 35*) which have regarded them as opposites. However, because previous investigations (*7*) have noted that opposite emotions may fluctuate somewhat independently of each other, we verify that our primary conclusions remain unchanged when we instead normalize using all loggable symptoms on the screen on which a mood appears – in essence, computing the probability that a person logged a given mood given that they looked at the screen on which it was an option to log. Our primary conclusions remain unchanged under this parameterization: the observed cycle trajectories remain consistent (with positive moods declining, and negative moods increasing, prior to period start), and the menstrual and daily cycles remain most prominent.

tracking the relevant data, we analyze only people who log the behavior at least once. We do not analyze the time at which behaviors are logged, since a person may log at 9 PM that they slept, had sex, or exercised that day, but it does not necessarily indicate they had slept, had sex, or exercised at 9 PM.

##### 1.2.5 Continuous dimensions: vital signs

For the three vital sign dimensions — basal body temperature (BBT), resting heart rate (RHR), and weight — our dataset does not contain the time at which the log was entered, so we apply no time filtering and do not analyze daily cycles. For RHR, we have less than a year of reliable data, and so we do not analyze seasonal cycles. We apply basic quality control filters to each vital sign to ensure data is reliable (since, in contrast to the binary mood and behavior logs, for vital signs people can enter implausible values). For weight, we filter out weights less than 50 or greater than 500 pounds; filter out people whose weight fluctuates by more than 50 pounds (since this indicates people who may be trying to lose weight, who may have different cyclic patterns, and may also indicate people who log unreliably) and filter out people with fewer than 5 logs. For RHR, which is automatically measured by a heart rate monitor, we filter out people with logs on fewer than 50 days and people whose RHR fluctuates by more than 50 BPM (all RHR observations are in a biologically plausible range, so we do not filter for a maximum or minimum value). For BBT data, we filter out BBTs below 90 degrees or greater than 110 degrees, people with fewer than 50 logs, and people with fewer than 5 unique values of BBT (since BBT is somewhat difficult to measure, and we observe that some people log only implausibly constant values of BBT). A small fraction of people have multiple readings on a single day; we average these together so that there is only a single observation for each person on each day.

#### 1.3 Decomposition into four cycles

We separate the overall signal into daily, weekly, seasonal, and menstrual cycles as follows. Because we are interested in *within*-individual variation, we first remove individual means, following previous literature (*7, 8*): that is, for each individual and each dimension, we subtract the individual's mean for that dimension, so each each individual has zero mean. We then run a linear regression of the observations $x$:

$$x_{yphwm} \sim \mathcal{N}(\alpha + \beta_y + \gamma_p + \delta_h + \eta_w + \kappa_m, \sigma^2)$$

where $y$ indexes year, $p$ indexes day relative to period, $h$ indexes hour, $w$ indexes weekday, $m$ indexes month, and all are encoded as categorical variables (with a distinct coefficient for each value). $y$ ranges from 2015 to 2017; $p$ ranges from -20 to 20; $h$ ranges from 0 to 23; $w$ is the seven days of the week; and $m$ ranges from 1 to 12. In all four-cycle decomposition plots (*e.g.*, Figure 2b), we extract the relevant coefficients and zero-mean them; the "Baseline" label on the plot indicates a coefficient of zero[4]. Errorbars are 95% confidence intervals. We plot only estimates for day relative to period from -14 to 14 because the average menstrual cycle is roughly 28-29 days long[5]. One advantage of our large dataset is that it allows us to precisely estimate day-specific coefficients for the menstrual cycle, rather than making potentially false parametric assumptions about how mood, behavior, and vital signs will change over the course of the cycle (*11*).

We define the *amplitude* of a cycle as the difference between the maximum and minimum

[4]Similarly, when we plot the daily signal in Figure 2a, we subtract off the mean signal across days, so the average signal value is zero; we label this as "Baseline" on the plot. To plot the red line in Figure 2a — the counterfactual world where menstrual cycle effects are observed rather than averaged out — we add the inferred menstrual cycle effect, as plotted in Figure 2b, to the observed signal (black line) in Figure 2a.

[5]Day -14 is not equivalent to day 14, since many women have cycles longer or shorter than the typical length of 28-29 days, so we would not necessarily expect the regression coefficients for day -14 and day 14 to be equivalent; some discontinuities at the graph boundaries are expected.

coefficient values for the cycle. We compute confidence intervals on this amplitude (Figure 2c) by bootstrapping replicate datasets, recomputing regressions and amplitudes for each replicate, and computing the 95% confidence interval of the bootstrapped amplitudes.

#### 1.4 Computation of near-period effects

For each dimension, we define the *near-period effect* by determining the week-long interval (beginning up to two weeks before period start) in which the dimension's mean value (as measured by the regression coefficients shown in Figures S1, S2, and S3) differs most dramatically from its overall mean (Table S4). Essentially, this captures the interval in which the cycle curve reaches its most significant peak or trough. All dimensions display their most pronounced peaks or troughs near period start, justifying our examination of a near-period effect; however, the exact timing of this peak or trough varies by dimension, as previous authors have also observed (*16*), justifying our use of a specific weeklong period for each dimension. For example, for sexual activity, the near-period interval begins at day -1 and ends at day 6 (where 0 denotes the period start date), and for BBT the near-period week begins at day -8 and ends at day -1.

After defining the near-period interval for each dimension, we define the *near-period effect* as the dimension's average value during the near-period interval minus the dimension's average value not during the interval. (As with the four-cycle analyses, we subtract each individual's mean prior to taking the averages). This is equivalent to performing a linear regression

$$x_p \sim \mathcal{N}(\alpha + \beta_p, \sigma^2)$$

where $p$ indicates that an observation occurs during the near-period interval. (We note that simply computing the amplitude of the near-period peak or trough, while intuitive for the population as a whole, would not allow us to perform a multiple regression, and thus to determine the effects of multiple covariates like age or country.) To compute country-specific period effects, we perform the regression:

$$x_{pc} \sim \mathcal{N}(\alpha + \beta_p + \gamma_c + \delta_{pc}, \sigma^2)$$

where $c$ denotes country, and the interaction term $\delta_{pc}$ allows period effects to differ by country. To confirm that our country-specific effects are robust to inclusion of other covariates (for example, age) we also fit models

$$x_{pca} \sim \mathcal{N}(\alpha + \beta_p + \gamma_c + \delta_{pc} + \eta_a + \kappa_{pa}, \sigma^2)$$

where $a$ denotes age group, and $\kappa_{pa}$ allows for age-specific period effects. Besides age, the additional covariates we include are behavior controls (if the person has ever logged consuming alcohol; logged consuming cigarettes; logged exercise; logged taking hormonal birth control; logged taking a birth control pill; or logged using an IUD); and app usage controls (number of symptom categories used; start year; and total symptoms logged). The goal of this analysis is to ensure that our country-specific estimates are not driven by other country-specific differences.

For each fitted model, we compute the country-specific period effect for each country $c$ by setting the country for all people to $c$ (keeping their other covariates the same) and computing the difference between their model-predicted value during the near-period interval and not during the near-period interval. This can be interpreted as the model's predicted near-period effect for each country if the distribution of all non-country covariates were that of the population as a whole[6].

[6]For computational tractability, given the large number of countries in the dataset, we compute this quantity on a random sample of 100,000 users; our confidence intervals (which are very small) are thus conservative. We compute 95% confidence intervals by resampling the entire dataset using bootstrapping, repeating the period effects estimation procedure on each bootstrapped replicate, and computing the 95% confidence interval of the bootstrapped period effects.

Our computations for age-specific period effects are analogous. In both cases, we find that inclusion of other covariates does not significantly change our age- or country-specific estimates. Our country-specific period effects are robust to inclusion of other covariates (Figure S7); similarly, we infer the same age trends regardless of what other covariates we include (Figure S8).

#### 1.5 Robustness checks

##### 1.5.1 Alternate four-cycle decomposition methods

We compare our linear regression four-cycle decomposition (as described in Section 1.3) to two other cycle decomposition methods: taking the means for each cycle separately, and fitting a mixed model.

- **Means by group**. Rather than fitting a linear regression, we fit a simpler model where, after subtracting the means for each individual, we simply compute the means for each day relative to period, hour, weekday, and month. This does not allow us to control for year or account for correlations between cycles (for example, if people tend to log when they are on their period only if it is a Sunday) but yields similar results to linear regression.
- **Mixed model**. We fit a mixed model which includes the same fixed effects as in the linear regression but, rather than removing the individual mean, we fit a random-effects intercept term for each individual. The motivation is that, while simply removing individual means is interpretable, scalable to large datasets, and used in previous studies (*7*), it can also potentially lead to misleading estimates of cyclic effects by incorrectly attributing cyclic variation to between-individual variation. For example, if each individual logs for only a week, removing individual means may attenuate seasonal cycle estimates. We use a linear mixed model even though some of the dimensions are binary because we want the fixed

effects coefficients to be interpretable and comparable to our estimates from the other two methods. We downsample the data for each dimension to a maximum of 100,000 individuals (randomly selected) for computational tractability.

Both methods yield very similar results to linear regression. The mixed model yields slightly larger estimates of cycle estimates (the estimated menstrual cycle amplitudes are 27% larger on average across all dimensions) so our estimates of cycle effects should be regarded as conservative. However, our main conclusions remain similar under the two alternative specifications: in particular, menstrual cycle amplitudes remain generally larger than those of the other three cycles. We therefore favor the linear regression model for its simplicity and because it scales to the entire dataset[7].

##### 1.5.2 Alternate parameterization of seasonal cycles

Because there are multiple ways of parameterizing seasonal cycles, we assess how our results vary under two alternate regression parameterizations: 1) replacing the month-of-year indicator variable $\kappa_m$ with a week-of-year indicator variable and 2) replacing the year indicator variable $\beta_y$ with a linear time trend. Our results remain similar under these alternate parameterizations. Replacing month-of-year with week-of-year slightly increases the amplitude of the seasonal cycle, as expected, due both to noise and to greater sensitivity to transient events like Christmas, but estimates for the seasonal cycle are similar and estimates for other cycles are nearly identical; we favor the month-of-year parameterization because it allows us to more robustly estimate the seasonal cycle for small subgroups of the population in our substratification robustness anal-

[7]The only case in which the models yield somewhat different results is for seasonal cycles in weight, for which the mixed model and linear regression model estimate qualitatively similar trends but the mixed model estimates a considerably larger amplitude (0.65 versus 0.24 lbs for the linear regression model). This discrepancy occurs because there is substantial change in the average weight of the population over time, likely caused by the expansion in the population using the app, and the mixed model attributes this to seasonal variation. Our estimates of seasonal changes in weight thus ought be regarded as conservative, and it is possible the true amplitude of the seasonal cycle effect is somewhat larger.

ysis (Figure S5). Replacing the year indicator variable $\beta_y$ with a linear time trend also produces generally similar results.

##### 1.5.3 Alternate parameterization of menstrual cycles

There are multiple ways of parameterizing the menstrual cycle; we explore three alternative parameterizations in addition to our primary parameterization (ranging from -20 – the nearest cycle start is 20 days in the future – to 20 – the nearest cycle start is 20 days in the past):

- **Day relative to last cycle start**: for every date on which a person logged, we compute the date of the *last* cycle start on or before the log date and take the difference in days. To ensure we have reliable information for cycle day, in this analysis we discard logs with more than 40 days since last cycle start.
- **Day relative to next cycle start**: for every date on which a person logged, we compute the date of the *next* cycle start on or after the log date and take the difference in days. We discard logs with more than 40 days until the next cycle start.
- **Fraction of the way through cycle**: because average cycle length varies across individuals, for each individual and each log, we compute the log's cycle day (as in our primary parameterization) and divide by the individual's average cycle length. For example, if the log occurred on cycle day 9, and the person's average cycle was 30 days long, the fraction of the way through the cycle $f$ would be $\frac{9}{30} = 0.3$. We analyze logs with $-0.5 \leq f \leq 0.5$.

Our estimated menstrual cycle amplitudes remain stable under these three alternate parameterizations of the menstrual cycle, confirming that our conclusions are robust to our parameterization of the menstrual cycle. Specifically, in all cases our estimated menstrual cycle amplitude is within 20% of our original estimate, and the average difference between the original and alternate parameterizations is 4.8%.

##### 1.5.4 Substratification robustness checks

The population using the app is likely non-representative; we therefore verify that the menstrual cycle remains prominent when we substratify the dataset by demographics (age group, country, and latitude), app usage variables (number of categories logged and number of symptoms logged), and whether individuals logged in at least twelve unique months and were included in the main analysis, to ensure that this filtering (Section 1.2) does not change conclusions (Figure S5). For each subgroup, we compute the amplitude of the menstrual, daily, weekly, and seasonal cycles for that subgroup alone. Unsurprisingly, we observe some variation in the cycle amplitudes across subgroups (variation across subgroups is, after all, the phenomenon that our age subgroup analysis highlights). However, importantly, the overall ordering of amplitudes remains largely stable across subgroups for each dimension and, in particular, the prominence of the menstrual cycle does not appear to be driven by any particular subgroup. This indicates that our central finding of the prominence of the menstrual cycle is unlikely to be driven by a non-representative subgroup.

#### 1.6 Reproducing previously known results

Here we describe our procedures for replicating previously known findings. The purpose of this analysis is to verify that our dataset is sufficiently reliable to reproduce previous results.

##### 1.6.1 Menstrual cycle lengths

The means and standard deviations of menstrual cycle lengths in our dataset match previous findings in Chiazze et al, 1968 (*36*) (Figure S9). Following those authors, we filter for cycles between 15 and 45 days in length and stratify by age group. Both the mean and standard deviations for each age group closely match the previous estimates; our data also recapitulates the slight decrease in means and standard deviations by age.

##### 1.6.2 Country-specific patterns

We compare our dataset's country-specific measures of happiness and weight to previous country-specific measures of happiness and weight. Our source for previous country-specific happiness data is the Gallup World Poll, which has been used in a previous study of happiness (*37*). Following those authors, we use Gallup's measures of real-time positive/negative experiences (as measured by experiences on the day before the survey) and overall life evaluation. We examine the correlations between Gallup's measures and our dataset's fraction of happy/sad logs which are happy across the 79 countries with Gallup data and at least 1,000 happy/sad logs from 100 unique individuals in our dataset. All correlations are statistically significant and have the expected sign: the correlation with Gallup's real-time positive experience index is $r = 0.51$ ($p < 0.001$); with Gallup's real-time negative experience index is $r = -0.38$ ($p < 0.001$); and with Gallup's overall life evaluation is $r = 0.47$ ($p < 0.001$).

For weight, we compare the average weight of individuals in our dataset who provide weight data to previous country-specific measures of weight and obesity. We study the 38 countries for which we have weight/obesity data from an external dataset and at least 1,000 weight logs from 100 individuals in our dataset. The average weight of individuals in each country (in our dataset) is significantly correlated with 2016 WHO estimates of the fraction of women over 18 who are overweight (*38*) (BMI $>$ 25) ($r = 0.53, p < 0.001$), the fraction of women over 18 who are obese (*39*) (BMI $>$ 30) ($r = 0.59, p < 0.001$), and the average weight of adults in that country (*40*) ($r = 0.51, p = 0.001$).

##### 1.6.3 Previously known cycles

In addition to the dimensions of mood, behavior, and vital signs considered in our primary analysis, we investigate whether our dataset displays expected seasonal, weekly, and menstrual cycles for additional dimensions (Figure S10). (We do not investigate additional dimensions in daily cycles because the dimensions for which we have the most reliable time information — mood symptoms — are included in our primary analysis.) As with all our analyses which include seasonal cycles, we filter for individuals in the Northern hemisphere.

For menstrual cycles, we examine the four physical pain symptoms the Clue app allows people to log – cramps, headache, ovulation pain, and tender breasts – since these are some of the best-studied and most commonly logged menstrual symptoms. Consistent with prior research, we observe in our dataset that cramps (*41, 42*), breast pain (*43*), and headache (*42*) are more commonly reported by individuals near period start. In contrast, individuals in our dataset are most likely to report ovulation pain at about day -14 (*i.e.*, 14 days before the start of their next period), consistent with prior findings of "Mittelschmerz" pain occurring near ovulation (*44*). (Ovulation pain also shows a slightly smaller peak at day 14 in our dataset — equivalent to day -14 for women with cycles near the typical length of 28 days.)

For weekly cycles, we examine patterns in alcohol consumption. Previous findings indicate that alcohol consumption increases on the weekend (*45, 46*); consistent with this, we observe that individuals are more likely to report attending parties with alcohol on the weekends, with logs of hangovers peaking on Sundays.

For seasonal cycles, we investigate allergies, cold/flu symptoms, and vacationing. Seasonal allergies have been previously found to peak in spring and summer because of higher pollen counts (*47–49*). Incidence of flu peaks in the winter (*50, 51*). Vacation is more commonly taken during the summer months and near the winter holidays (*52*). Our dataset displays patterns consistent with all these prior findings.

#### 1.7 Generalizability of findings

Since women who use menstrual cycle tracking apps likely differ from the general population, it is natural to ask how our findings generalize. While a population of millions of women is arguably large enough to be worth studying in and of itself even if it is somewhat non-representative, three lines of evidence support the generalizability of our findings.

1. Menstrual tracking apps are increasingly widely used (*24*), offering a data source which is plausibly more representative than those used in previous studies of the menstrual cycle, which have used small and non-representative populations (*20*).

2. Previous studies of the dataset have found that it replicates known biology — for example, menstrual cycle lengths (*14*) and premenstrual symptoms (*15*), (*16*). Our present analysis similarly produces numerous findings consistent with previous studies: a late-night shift towards negative mood (*7*) (*53*); estimates of menstrual BBT, RHR, and weight cycle amplitudes (*23*) (*32, 54, 55*); decreases in exercise on the weekends and during the winter (*56, 57*); and decreases in sexual activity immediately after the period begins and during the weekdays (*58*). In addition to our main analysis, we replicate a number of other previous results on our dataset (Section 1.6). These replications suggest that our dataset is sufficiently reliable to reproduce previous results.

3. Our results remain stable and the menstrual cycle remains prominent when we substratify the dataset by demographics and app usage variables (Section 1.5, Figure S5), indicating that our results are unlikely to be driven, for example, by very young women or by women who use the app more frequently because they have particularly severe menstrual symptoms.

#### 1.8 Reliability of self-reported data

Two lines of evidence increase our faith that the data is reliably logged. First, a review of more than a hundred menstrual cycle tracking apps rated the Clue app as the best on the basis of accuracy, comprehensiveness, and functionality (*13*), increasing the likelihood that its data is reliable. Second, we apply several quality control filters (Section 1.2) to ensure the accuracy of logged data: for mood and behavior dimensions, we filter for logs that are *prospectively* recorded (logged on the same day on which they occur), since prospective logging is more reliable (*7*). For vital sign dimensions, we filter for individuals who log frequently and remove biologically implausible values. For mood dimensions, one concern is that because the app interface renders the menstrual cycle more salient, it primes people to report subjective period-related mood changes. This is an unavoidable concern in studies of menstrual mood changes; further, if menstrual cycle apps are priming large fractions of the female population to experience negative mood near their periods, that is itself a phenomenon worth studying. The more objectively recorded behavior and vital sign dimensions should be unaffected by this phenomenon.

#### 2 Supplementary Tables and Figures

| | Long term loggers | | | |
|---|---|---|---|---|
| | **Mean** | **Median** | **5th percentile** | **95th percentile** |
| **Age** | 22.5 | 19.9 | 14.0 | 38.7 |
| **Days using app** | 541 | 538 | 348 | 727 |
| **Total logs entered** | 263 | 109 | 25 | 1004 |

| | All people using app | | | |
|---|---|---|---|---|
| | **Mean** | **Median** | **5th percentile** | **95th percentile** |
| **Age** | 21.2 | 18.5 | 13.4 | 37.2 |
| **Days using app** | 221 | 166 | 1 | 629 |
| **Total logs entered** | 105 | 36 | 3 | 411 |

Table S1: Summary statistics for long-term loggers (top), whom we use in our primary analyses, and all people using the app (bottom), whom we analyze as a robustness check. Age is computed using only people for whom age data is available. "Days using app" indicates the number of days between the first date a person entered a log on the app and the last date they entered a log on the app.

| Category | Specific features | % of logs |
|---|---|---|
| emotion | PMS, happy, sad, sensitive | 11% |
| period | heavy, light, medium, spotting | 10% |
| energy | energized, exhausted, high energy, low energy | 9% |
| sleep | 0-3 hrs, 3-6 hrs, 6-9 hrs, 9 hrs | 8% |
| pain | cramps, headache, ovulation pain, tender breasts | 8% |
| skin | acne, dry, good, oily | 6% |
| mental | calm, distracted, focused, stressed | 6% |
| motivation | motivated, productive, unmotivated, unproductive | 6% |
| craving | carbs, chocolate, salty, sweet | 6% |
| social | conflict, sociable, supportive, withdrawn | 5% |
| digestion | bloated, gassy, great digestion, nauseated | 4% |
| hair | bad, dry, good, oily | 4% |
| poop | constipated, diarrhea, great, normal | 3% |
| collection method | menstrual cup, pad, panty liner, tampon | 3% |
| sex | high sex drive, protected, unprotected, withdrawal | 2% |
| pill birth control | double, late, missed, taken | 2% |
| fluid | atypical, creamy, egg white, sticky | 2% |
| vital signs | BBT, heart rate, weight | 1% |
| exercise | biking, running, swimming, yoga | 1% |
| medication | antibiotic, antihistamine, cold/flu medication, pain | $< 1\%$ |
| ailment | allergy, cold/flu ailment, fever, injury | $< 1\%$ |
| party | big night party, cigarettes, drinks party, hangover | $< 1\%$ |
| appointment | date, doctor, ob/gyn, vacation | $< 1\%$ |
| test | ovulation negative, ovulation positive, pregnancy negative, pregnancy positive | $< 1\%$ |
| iud | inserted, removed, thread checked | $< 1\%$ |
| ring birth control | removed, removed late, replaced, replaced late | $< 1\%$ |
| injection birth control | administered | $< 1\%$ |
| patch birth control | removed, removed late, replaced, replaced late | $< 1\%$ |

Table S2: All features logged in the dataset. Features are organized into categories by Clue: for example, “PMS”, “happy”, “sad”, and “sensitive” features are all *emotion* features. Categories are sorted by how frequently they are logged in the dataset.

| Category | Dimension | Mean (LTLs) | Mean (overall) | $n_{\text{obs}}$ (LTLs) | $n_{\text{obs}}$ (overall) | $n_{\text{people}}$ (LTLs) | $n_{\text{people}}$ (overall) |
|---|---|---|---|---|---|---|---|
| Vital sign | RHR (BPM) | 65.3 | 65.3 | 1,485,816 | 1,849,409 | 8,939 | 13,070 |
| Vital sign | Weight (LB) | 135.2 | 134.7 | 1,243,469 | 3,074,289 | 47,540 | 184,808 |
| Vital sign | BBT (deg F) | 97.6 | 97.6 | 540,463 | 974,982 | 3,533 | 9,033 |
| Mood | happy vs sensitive | 58.8% | 58.8% | 10,075,794 | 32,408,134 | 415,690 | 2,779,666 |
| Mood | happy vs sad | 75.4% | 75.7% | 7,858,798 | 25,178,082 | 401,418 | 2,597,554 |
| Mood | calm vs stressed | 62.2% | 62.1% | 4,666,414 | 13,475,101 | 244,810 | 1,318,281 |
| Mood | motivated vs unmotivated | 51.0% | 50.7% | 4,529,869 | 12,996,012 | 245,866 | 1,287,296 |
| Mood | social vs withdrawn | 63.0% | 64.0% | 4,486,865 | 13,245,834 | 212,010 | 1,186,117 |
| Mood | productive vs unproductive | 53.7% | 49.7% | 4,171,684 | 11,661,326 | 235,765 | 1,220,912 |
| Mood | focused vs distracted | 49.2% | 47.5% | 3,972,562 | 11,772,608 | 236,967 | 1,267,988 |
| Mood | energized vs exhausted | 36.2% | 38.6% | 2,878,582 | 9,653,503 | 327,058 | 1,903,751 |
| Mood | supportive vs conflict | 64.2% | 61.7% | 2,094,079 | 6,533,528 | 179,270 | 958,396 |
| Behavior | had sex | 13.4% | 17.2% | 17,691,362 | 38,767,857 | 249,356 | 1,196,621 |
| Behavior | exercised | 13.0% | 17.5% | 11,643,466 | 25,722,601 | 163,175 | 778,307 |
| Behavior | slept $>$ 6 hours | 69.3% | 67.7% | 10,626,280 | 33,994,360 | 403,175 | 2,748,894 |
| **All combined** | - | - | - | **87,965,503** | **241,307,626** | **499,330** | **3,322,769** |

Table S3: The fifteen dimensions included in the cycles analysis. The “mean” columns provide the average value of the dimension: for example, 58.8% of happy/sensitive logs are happy. $n_{\text{obs}}$ provides the number of observations for that dimension and $n_{\text{people}}$ provides the number of people who logged that dimension. We report statistics for both long-term loggers (LTLs), whom we use in our primary analysis, and the overall dataset, which we also analyze as a robustness check.

| Category | Dimension | Start day | End day |
|---|---|---|---|
| Behavior | exercised | 0 | 7 |
| Behavior | had sex | -1 | 6 |
| Behavior | slept $>$ 6 hours | -10 | -3 |
| Mood | happy versus sad | -5 | 2 |
| Mood | happy versus sensitive | -5 | 2 |
| Mood | energized versus exhausted | -5 | 2 |
| Mood | calm versus stressed | -6 | 1 |
| Mood | focused versus distracted | -5 | 2 |
| Mood | motivated versus unmotivated | -5 | 2 |
| Mood | productive versus unproductive | -5 | 2 |
| Mood | social versus withdrawn | -5 | 2 |
| Mood | supportive versus conflict | -7 | 0 |
| Vital sign | BBT (deg F) | -8 | -1 |
| Vital sign | RHR (BPM) | -7 | 0 |
| Vital sign | Weight (LB) | -4 | 3 |

Table S4: Start and end days for the near-period interval for each dimension. Time intervals exclude the end day. Day 0 is the day of period start.

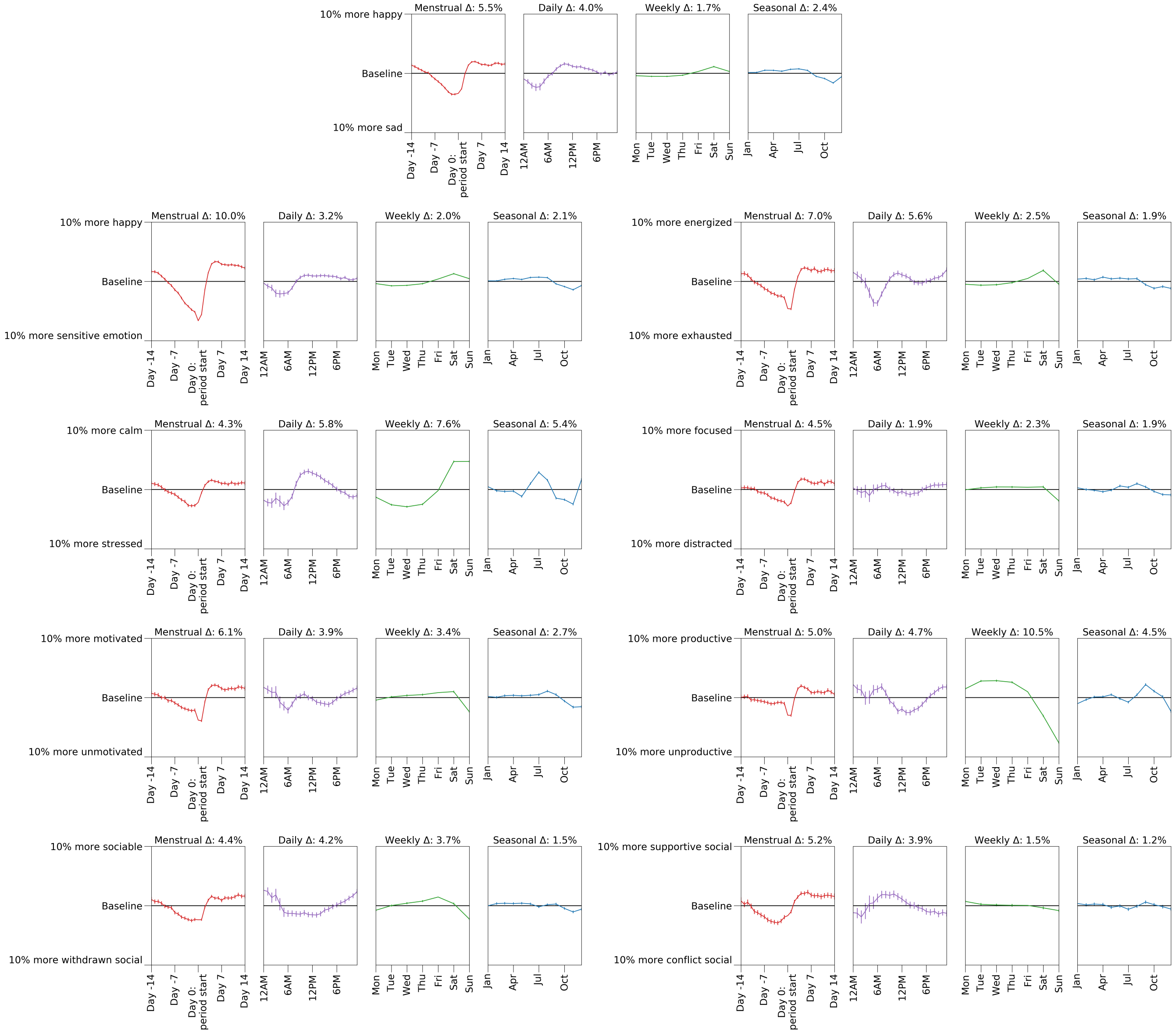

Figure S1: Four cycle plots for all mood dimensions as inferred by linear regression (Methods Section 1.3). Δ indicates the amplitude of each cycle (the difference between the maximum and minimum values).

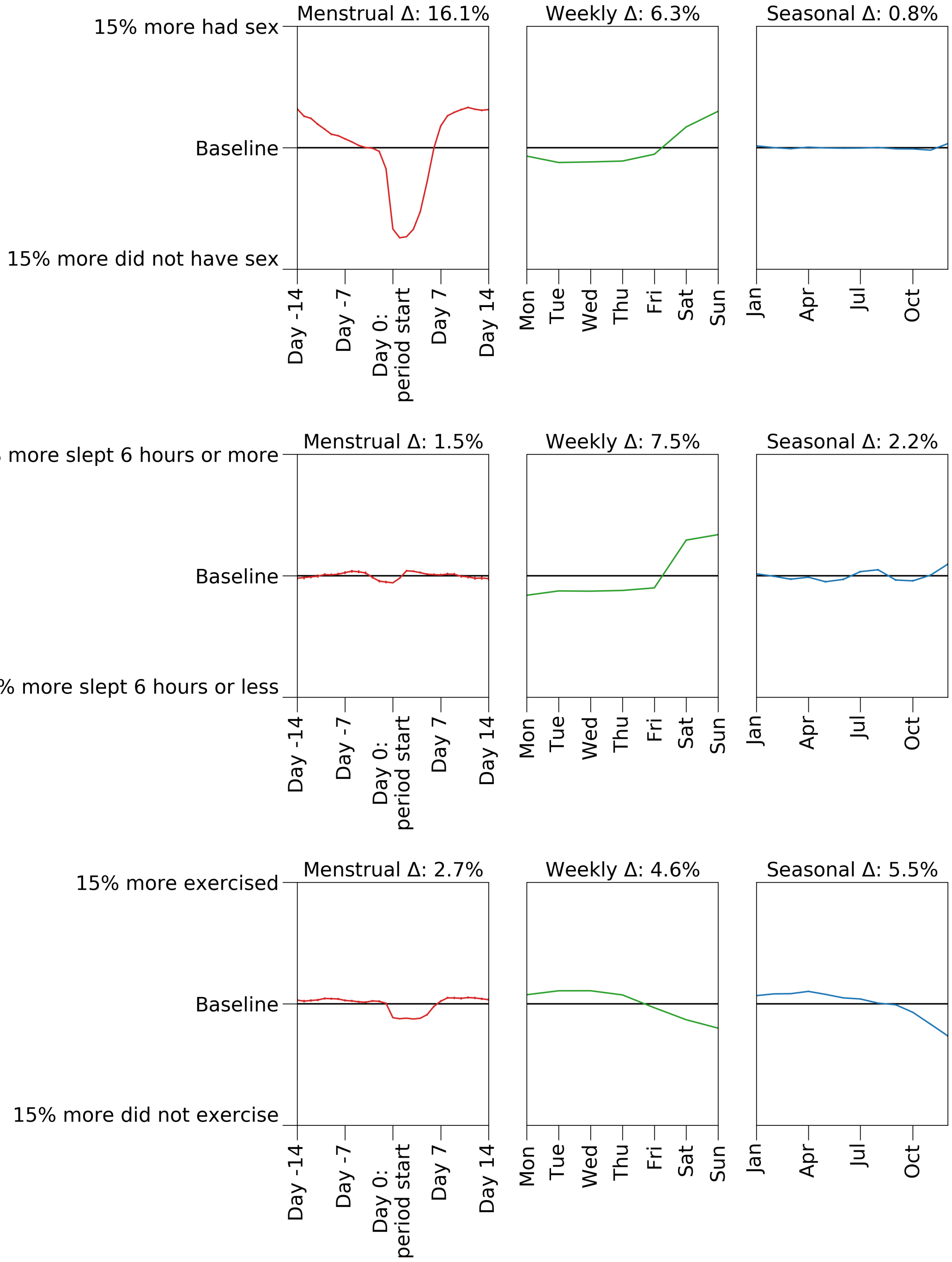

Figure S2: Four cycle plots for all behavior dimensions. No hourly information is available, so the daily cycle is omitted.

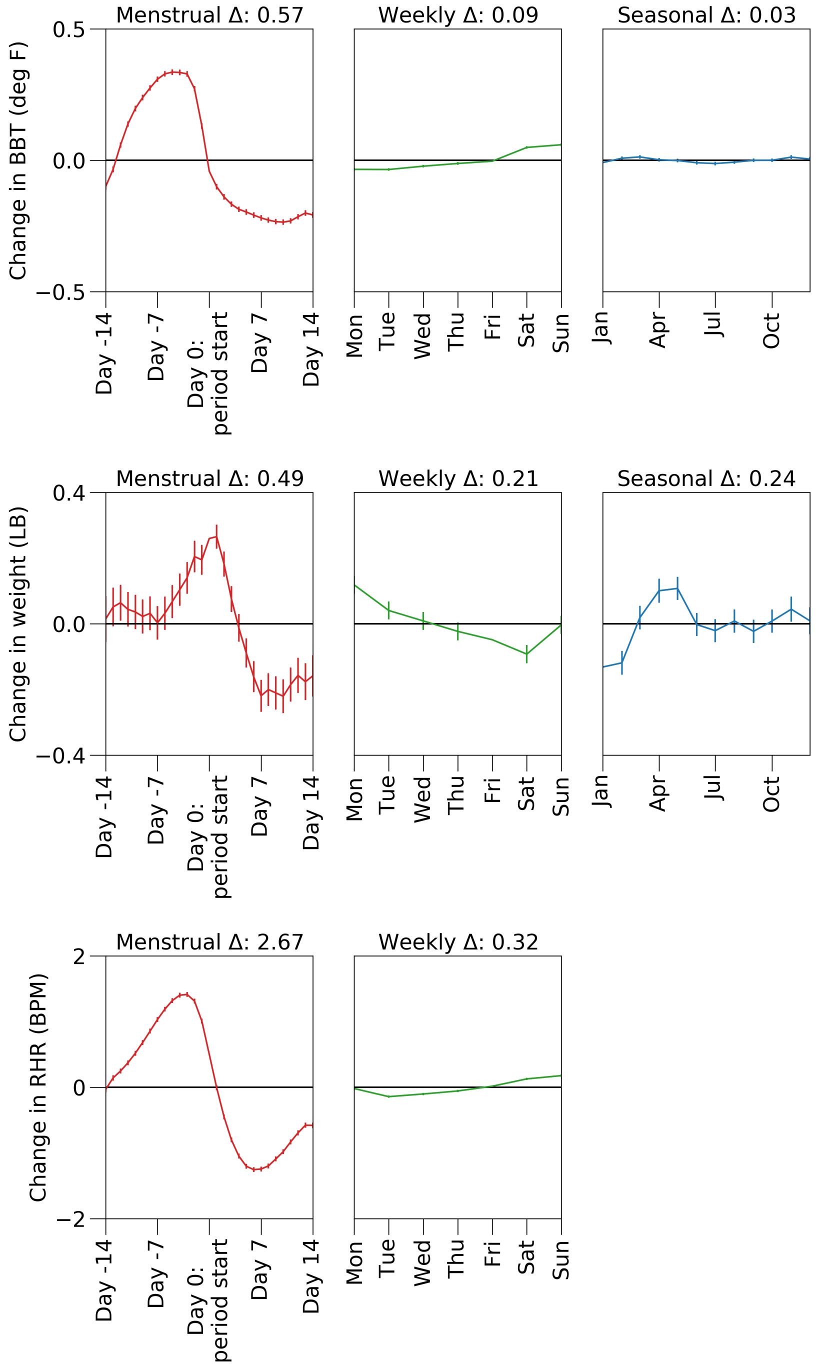

Figure S3: Four cycle plots for all vital sign dimensions: basal body temperature (BBT), weight, and resting heart rate (RHR). No hourly information is available, so the daily cycle is omitted. For RHR, the seasonal cycle is omitted because there is less than a year of data.

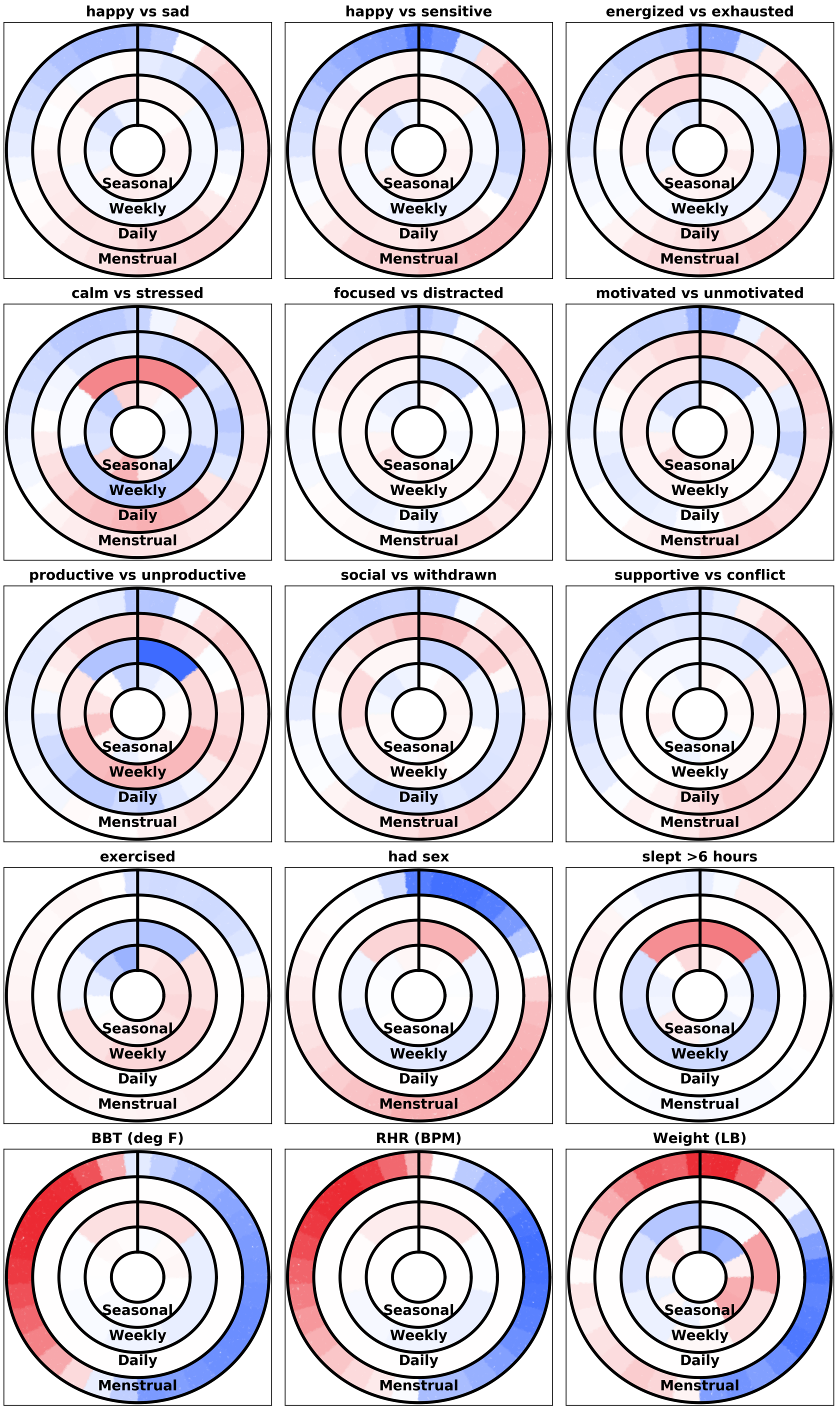

Figure S4: An alternate visualization of the four simultaneous cycles; this plot contains the same data as the four cycle plots above. Each subplot represents one dimension. Each concentric circle represents one cycle: from the outside progressing inward, the circles represent the menstrual, daily, weekly, and seasonal cycles. Cycles progress clockwise around the circles. The vertical black line denotes the start of the period for the menstrual cycle, midnight for the daily cycle, the middle of the weekend for the weekly cycle, and the start of the year for the seasonal cycle. More intense color changes indicate cycles with larger amplitudes, with red indicating positive changes, blue indicating negative changes, and white indicating zero change. Cycles lacking data for a dimension are shown in white.

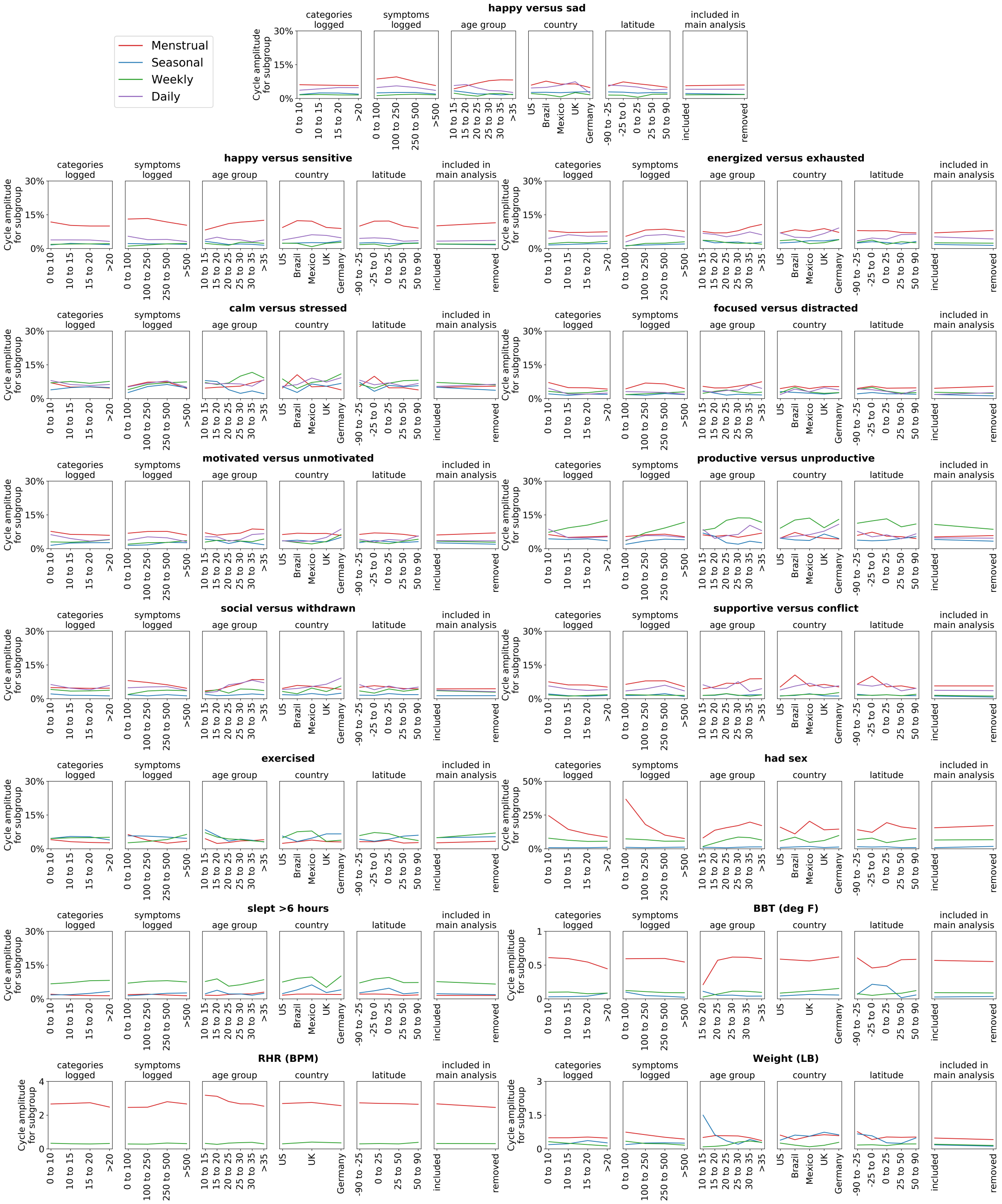

Figure S5: The menstrual cycle remains prominent across subsets of the population. Each set of subplots shows how cycle amplitudes for one dimension of mood, behavior, or vital signs vary over subsets of the population. The vertical axis plots the cycle amplitude; the horizontal indicates the subset of the population. For each dimension, the first two subplots subset by app usage patterns; the next three subplots subset by demographics; the final subplot subsets by whether individuals logged in at least twelve unique months and were included in the main analysis, to ensure that filtering does not change conclusions. While cycle amplitudes fluctuate across subsets of the population (as expected) the vertical ordering of the lines remains generally stable and, for dimensions in which the menstrual cycle is prominent, its prominence is not driven by a single subgroup. To avoid noisy estimates, only subsets which have at least 1,000 observations and 100 individuals are shown.

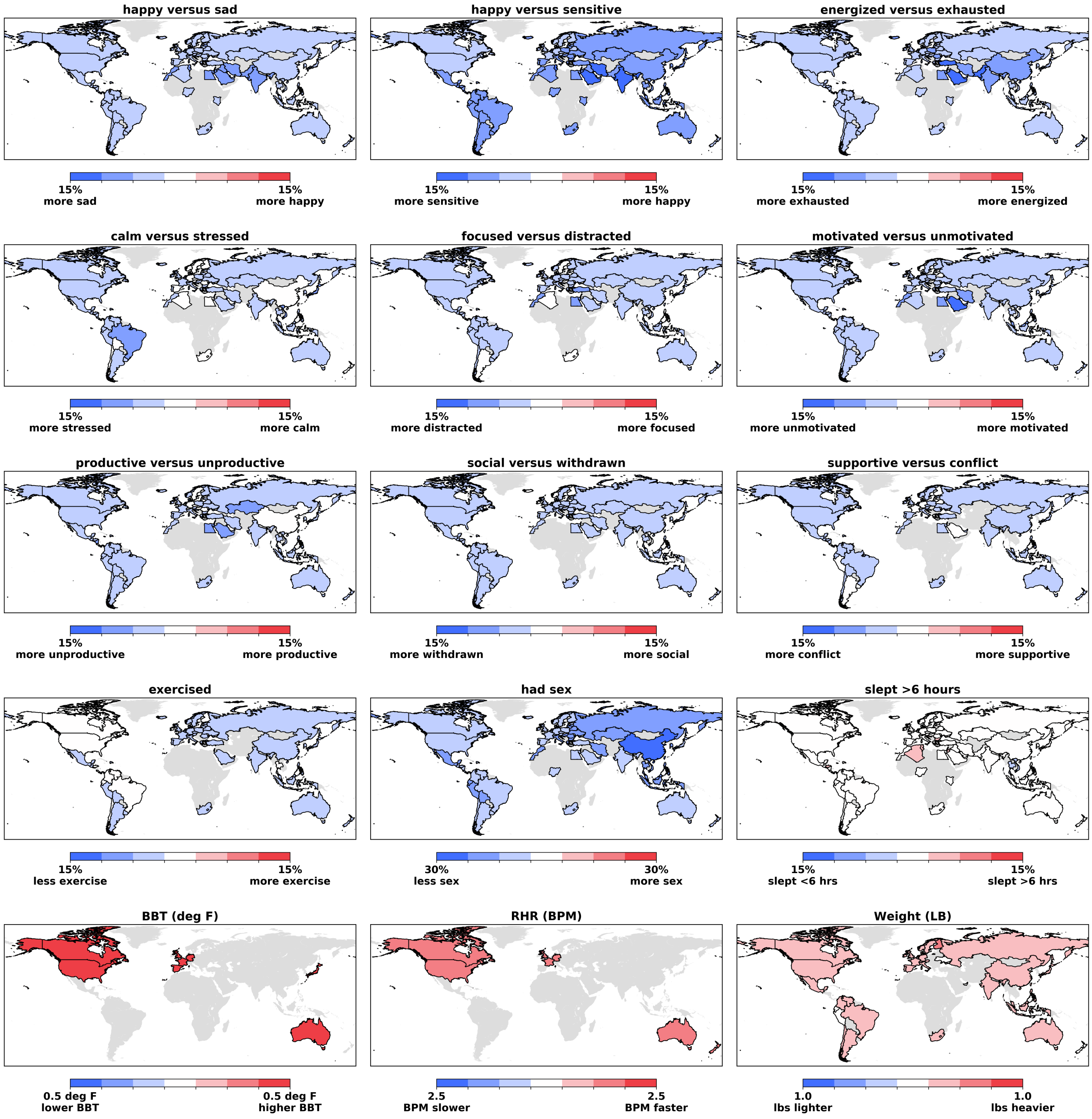

Figure S6: Maps of near-period effects for all cycle dimensions illustrate that effects remain largely directionally consistent across countries. Color bins are equally sized, with the central white bin centered on no change from baseline. Countries with fewer than 1,000 observations and 100 individuals for a dimension are shown in grey; for BBT and RHR, fewer countries have sufficient data but effects remain consistent.

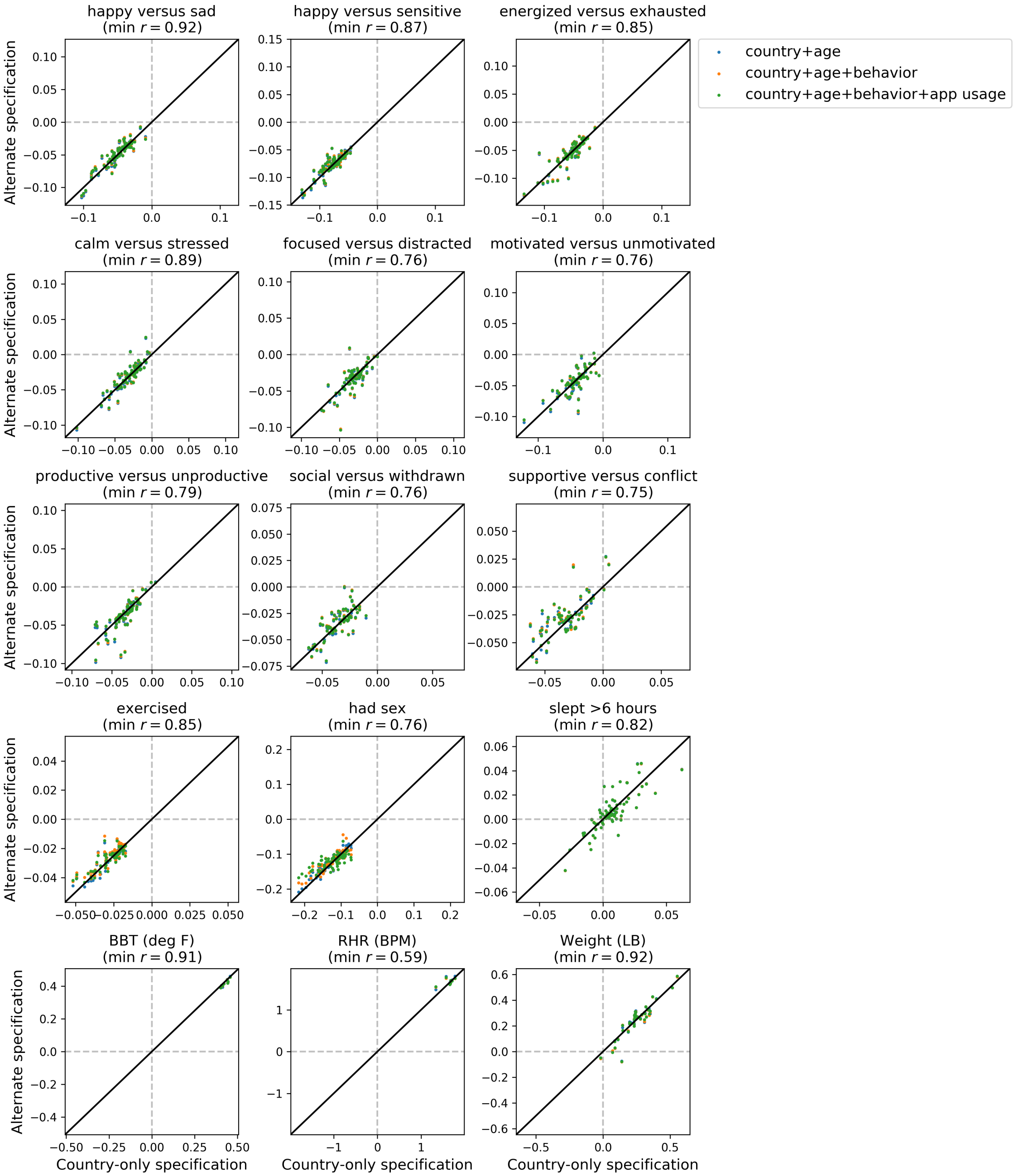

Figure S7: Near-period effects by country are robust to inclusion of other covariates. Each point represents one country (filtering for countries with at least 1,000 observations). The horizontal axis plots the estimated near-period effect using the simplest regression specification, with only country as a covariate; the vertical axis plots the estimated near-period effect using all other regression specifications (legend). Estimates are very similar, lying near the black diagonal line which indicates equality; minimum correlation is reported in the title of each plot.

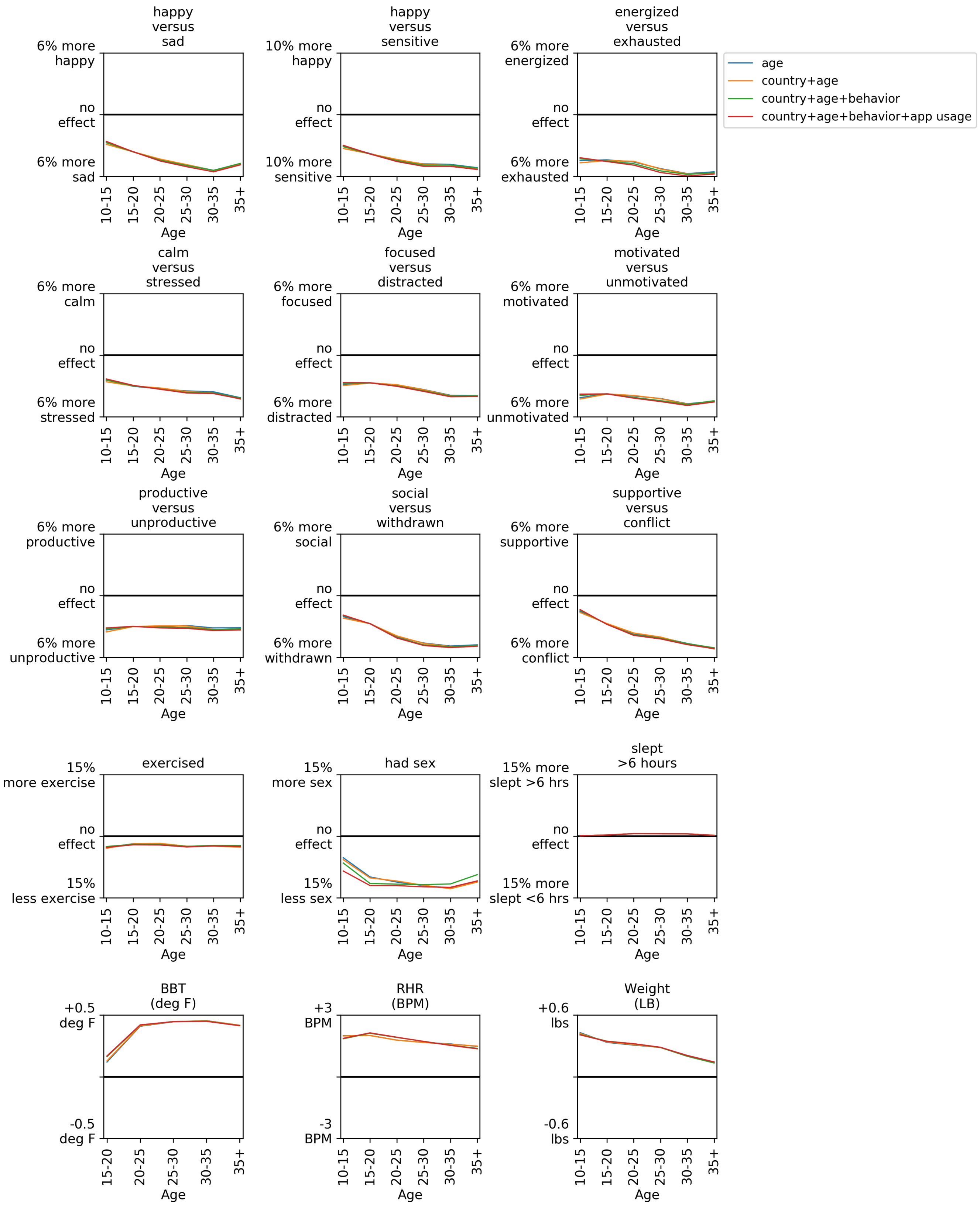

Figure S8: Near-period effects (vertical axis) by age group (horizontal axis); age trends are robust to inclusion of country, behavior, and app usage covariates. Different regression specifications are shown as different colored lines (legend), and in most cases the lines are too closely overlapping to be seen. All age groups have at least 1,000 observations and 100 individuals.

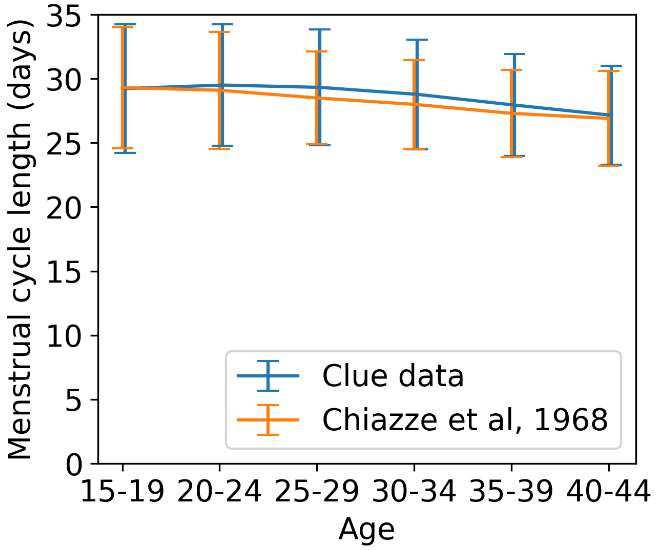

Figure S9: The mean and standard deviation of the menstrual cycle length distribution in our dataset is similar to that in Chiazze et al, 1968. Following their analysis, we filter for cycles between 15 and 45 days in length and stratify by age groups. Errorbars denote the standard deviation for menstrual cycles for each age group. Both the means and standard deviations are similar for each age group.

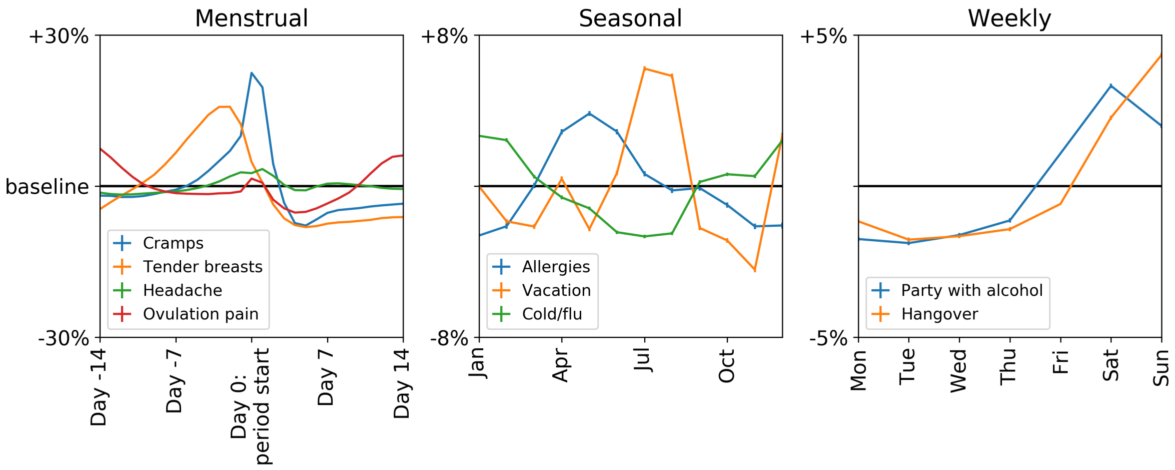

Figure S10: The dataset recapitulates previously known menstrual, seasonal, and weekly cycles.
